# Zero-shot antibody design in a 24-well plate

**DOI:** 10.1101/2025.07.05.663018

**Authors:** Chai Discovery Team, Jacques Boitreaud, Jack Dent, Danny Geisz, Matthew McPartlon, Joshua Meier, Zhuoran Qiao, Alex Rogozhnikov, Nathan Rollins, Paul Wollenhaupt, Kevin Wu

## Abstract

Despite breakthroughs in protein design enabled by artificial intelligence, reliably designing functional antibodies from scratch has remained an elusive challenge. Recent works show promise but still require large-scale experimental screening of thousands to millions of designs to reliably identify hits. In this work, we introduce Chai-2, a multimodal generative model that achieves a 16% hit rate in fully *de novo* antibody design, representing an over 100-fold improvement compared to previous computational methods. We prompt Chai-2 to design *≤* 20 antibodies or nanobodies to 52 diverse targets, completing the workflow from AI design to wet-lab validation in under two weeks. Crucially, none of these targets have a preexisting antibody or nanobody binder in the Protein Data Bank. Remarkably, in just a single round of experimental testing, we find at least one successful hit for 50% of targets, often with strong affinities and favorable drug-like profiles. Beyond antibody design, Chai-2 achieves a 68% wet-lab success rate in miniprotein design – routinely yielding picomolar binders. The high success rate of Chai-2 enables rapid experimental validation and characterization of novel antibodies in under two weeks, paving the way toward a new era of rapid and precise atomic-level molecular engineering.

*52 antigens targeted by Chai-2. Blue boxes indicate targets with at least one successful binder out of ≤20 assayed designs, representing 50% of the tested targets*.
**Date:** June 30, 2025

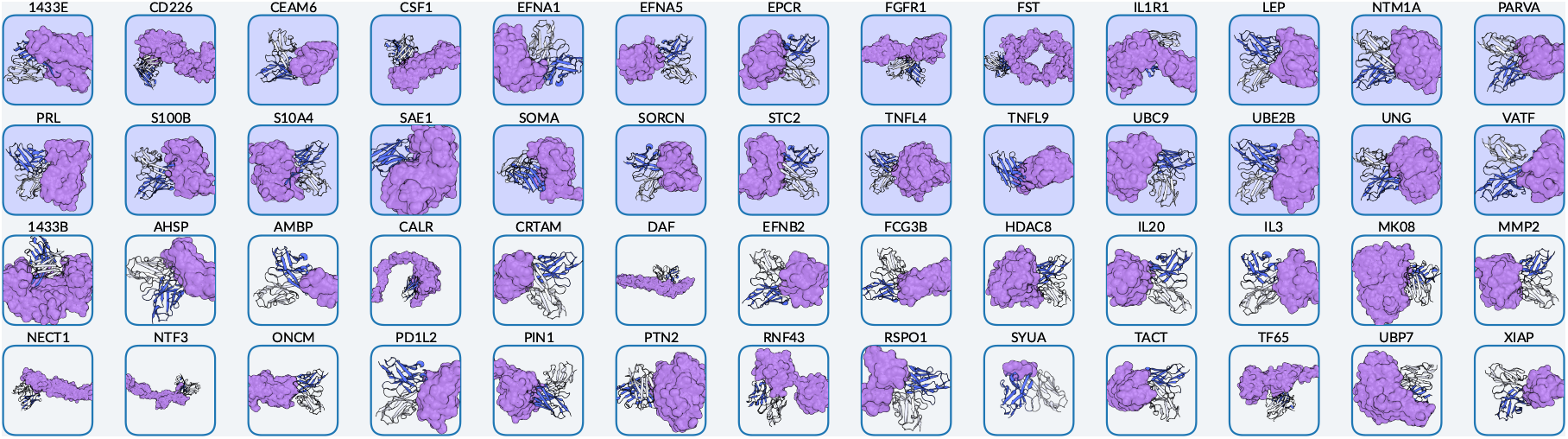

## 1 Introduction

Antibodies emerged over 500 million years ago to recognize molecular targets selectively and with high precision [1]. These properties, coupled with often favorable biophysical and immunogenic characteristics, have made antibodies highly desirable as therapeutics.

Today, antibodies are predominantly discovered by animal immunization campaigns or the screening of large immune repertoires or synthetic libraries [2]. These methods have helped facilitate substantial progress in the discovery process, resulting in the rapid growth of antibodies as a key therapeutic modality. By 2022, monoclonal antibodies constituted more than half of all biopharmaceutical approvals in the US and Europe, up from only 20% in the early 2000’s. [3]

Despite their success, traditional discovery pipelines remain resource-intensive and time-consuming, and frequently yield leads that demand months of downstream affinity or developability optimization. Moreover, these methods offer limited control of the desired binding site and often fail to hit hard targets or challenging-to-reach epitopes altogether.

These challenges have inspired significant research into computational methods for antibody discovery, which promise to deliver high-quality candidates at scale. Machine learning methods have driven significant advancements in molecular structure prediction [4, 5, 6, 7, 8, 9, 10, 11, 12, 13, 14], and more recently in protein design [15, 16, 17, 18, 19, 20, 21]. Although several groups have reported *de novo* antibody generation pipelines [22, 23, 24, 25, 26], none have demonstrated broad generalization to many targets and their hit rates rarely exceed 0.1% in the laboratory. As a result, these methods still rely on high-throughput experimental screening methods, undermining many of the principal benefits that they are intended to address.

Our main contributions are:

- We introduce **Chai-2**, which (to our knowledge) is the first fully *de novo* generation platform to design antibody binders with success rates high enough to reliably skip high-throughput screening.
- We successfully design diverse classes of binders—including miniproteins, antibody variable heavy chains (VHHs) and single chain variable fragments (scFvs)—across more than 50 targets, achieving state-of-the-art experimental success rates in all categories.
- We design and test *≤*20 antibodies or nanobodies per target, and found at least one experimentally confirmed *de novo* binder to 26 out of 52 novel targets.
- Designed antibodies are novel, diverse, and exhibit favorable developability profiles *in-silico*.
- We show Chai-2 can further optimize designs for specific therapeutic requirements such as species cross-reactivity.

The double-digit success rates we observe suggest that our method could enable discovery in a single 24-well plate, reducing experimental timelines to the order of weeks, and tightening the design–validation feedback loop. The high success rates across targets, with successful binders to roughly half of the antigens we tested, suggests this method could become broadly deployed. We believe our approach has the potential to reshape the de facto strategies of biologics lead discovery, as well as to address targets that have been challenging for traditional methods.

## 2 Methods

### 2.1 An all-atom foundation model for general purpose protein design

Chai-2 incorporates numerous advancements in all-atom generative modeling. For example, the Chai-2 folding module predicts antibody-antigen complexes with experimental accuracy twice as often as our previous Chai-1 model (Figures S1 and S2). Chai-2 generates candidate binders for any specified binding-site defined by just a few residues (Figure 1a) entirely “zero-shot” and without requiring a known starting binder. In addition, Chai-2 can generate sequences in a variety of modalities – including scFv antibodies, VHH domains, or minibinders – and can be prompted with multiple targets simultaneously, yielding proteins with tailored cross-reactivity and selectivity. Crucially, all of these capabilities are achieved without per-target tuning, demonstrating Chai-2’s scalability across diverse applications.

**Figure 1.**
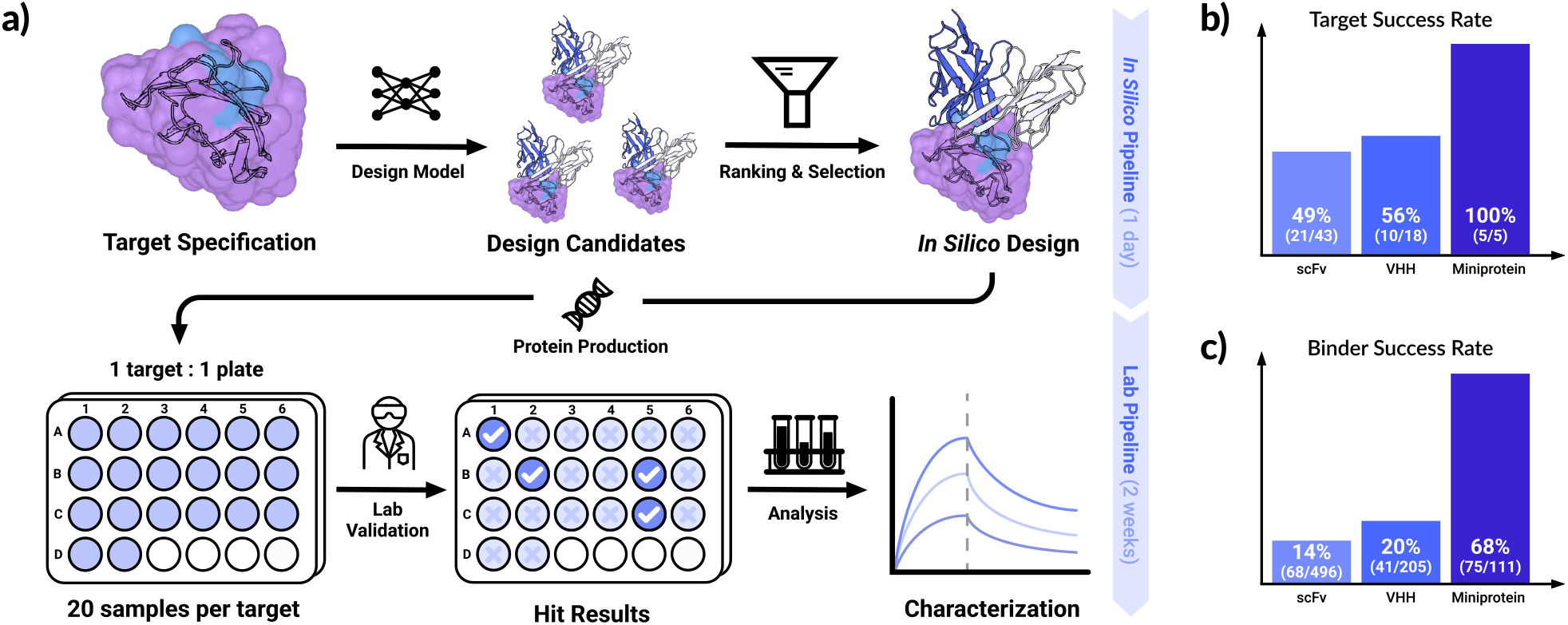
The Chai-2 model series reliably enables high-affinity protein binder and antibody design with state-of-the-art success rates. **(a)** The Chai-2 pipeline. Starting from a defined target structure and a list of epitope residues, the design model generates a sequence and all-atom structure that specifically interacts with the epitope. An *in-silico* ranking and selection model is employed to prioritize sampled designs. The top selected designs are directly advanced to fast experimental characterization that typically fits in a small-scale plate format such as single-concentration biochemical assays; the identified binders are further analyzed to determine quantitative binding affinity. Only two weeks are required to go from epitope to experimentally validated binders. **(b)** The percentage of targets per design modality (miniproteins, scFv, VHHs) where designs yield at least one experimentally validated binding design. **(c)** The percentage of total designs per modality exhibiting binding. Although it is difficult to directly compare lab success rates to previous work due to the limited number of targets previously reported, we note that the best reported success rates for *de novo* antibody design are typically lower than 0.1%.

### 2.2 Broad experimental validation of design capabilities on unbiased targets

We evaluate Chai-2 on miniprotein and antibody/nanobody design tasks. We select a separate panel of targets for each task; (1) a benchmark set from prior miniprotein studies to facilitate comparison, and (2) set of 52 novel antigens with no known antibodies in the Protein Data Bank (PDB, [27]). We provide further details for these sets through the remainder of this section. Additional information regarding target selection can be found in Section S1.

For miniprotein design, we test five targets previously studied in Cao et al. [28], AlphaProteo [20] or RFD-iffusion [18] (Table S1). Prior *in silico* methods have successfully designed miniproteins to four of the five targets: Interleukin-7 Receptor*α* (IL-7R*α*), Insulin receptor (InsulinR), Programmed Death-Ligand 1 (PD-L1), and platelet-derived growth factor receptor *β* (PDGFR*β*). As an additional challenge, we also test TNF*α*, a therapeutically important [29, 30] target estimated to be within the top 1% of difficulty among all potential targets in the PDB for *in silico* design [20]. The functional assembly of TNF*α* is a symmetric homotrimer that binds its native receptors (TNFR1 and TNFR2) through an interface between two subunit chains. The highly flat and polar nature of this binding site makes TNF*α* particularly challenging for computational protein design. To our knowledge, no prior computational work has *de novo* designed a protein binding TNF*α*.

To rigorously profile Chai-2’s ability to design antibodies against fully novel targets, we directly select proteins that were in stock in vendor catalogs, while excluding all proteins homologous to any antigen in the structural antibody database (SAbDab) [31] released prior to our training cutoff at 70% sequence identity and 80% coverage (see Section S1.2). Not only does this remove targets that have known antibodies themselves, but this also removes targets *similar* to chains that have known antibodies. This creates a challenging, unbiased set of novel targets that should assess Chai-2’s ability to design antibodies against arbitrary, unseen protein targets. To efficiently select meaningful epitopes, we further filter to examples with a known (non-antibody) binding partner. We use the resulting target protein and up to four residues on its native binding interface as a prompt to Chai-2. These targets represent a diverse set of proteins, from various cellular contexts and disease implications.

### 2.3 Experimental validation of candidate binders

For each target, we select up to 20 generated designs for experimental validation. We hypothesized that improvements in the quality of our models would allow us to avoid high-throughput experimental screening, and instead jump directly to individual characterization of each design.

We note that we **did not perform successive rounds of wet lab experimentation** and this was our first attempt ever at generating a binder for nearly every target tested. As such, results reflect the blind performance of running Chai-2 *once* in a new setting. We assess the binding strength of our designs by bio-layer interferometry (BLI) and we classified positive binding hits as designs with binding-positive curve signature while requiring that the signal is both greater than 0.1 nm above background and greater than 300% of background. We also perform additional validation to characterize other aspects of our designs. See Supplementary Information Section S5 for additional details.

## 3 Results

### 3.1 State of the art success rates in *de novo* miniprotein design

Computational miniprotein design has been thoroughly studied in the literature [28, 18, 20]. We therefore used this task as an initial benchmark. Using the five aforementioned targets (Table S1), we experimentally validate 20 designs for each from an initial prototype Chai-1d model, and 20-25 designs from Chai-2: PD-L1 (25); IL-7Ra (23); InsulinR (20); TNF*α* (22); PDGFR*β* (21)^1^. Experimental hit rates are shown on Figure 2a, juxtaposed with hit rates reported by prior works [18, 20]. For every target evaluated, Chai-2 achieves at least a three fold improvement in experimental hit rate compared to the next-best method and is even able to discover (to our knowledge) the first computationally designed hit against TNF*α*. A distribution of binding affinity measurements (*K*_D_) are shown for all discovered binders in Figure 2b. We observe picomolar *K*_D_values for IL7Ra, PD-L1, PDGFR*β*, and Insulin R, and low-nanomolar affinities for TNF*α*. We show examples of designed binders for all targets along with their binding affinities (*K*_D_) in Figure 2d. Chai-2 generates a variety of secondary structure elements, including a single-digit nanomolar PDGFR*β* binder composed almost entirely of beta sheets.

**Figure 2.**
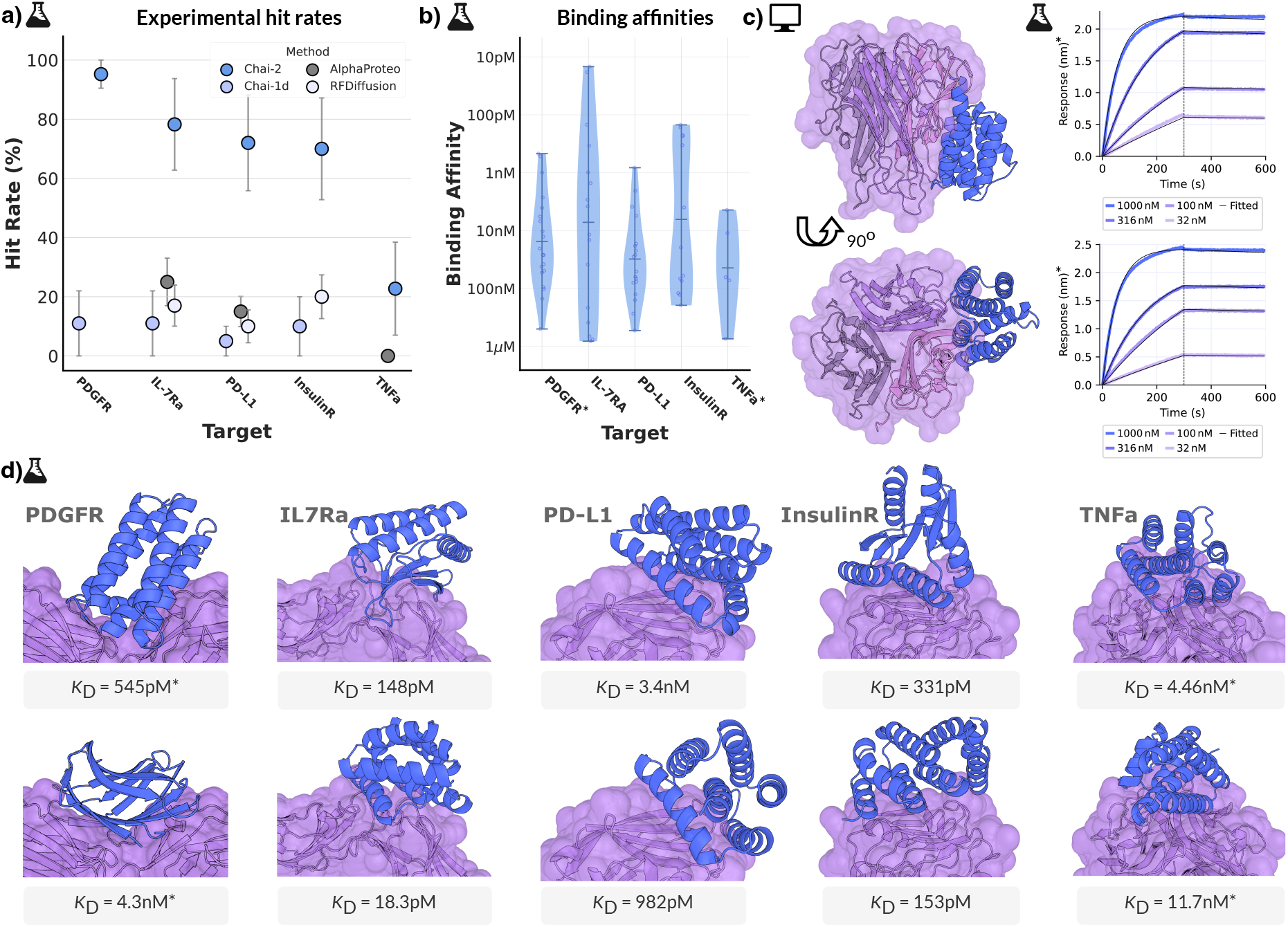
Chai-2 achieves state of the art performance on minibinder design. **(a)** Experimental hit rates of miniprotein generations from a prototype design model Chai-1d model (light blue) and our final Chai-2 model (dark blue). We compare to experimental success rates reported in the literature (shades of gray) [18, 20]. Error bars represent 95% confidence interval under a binomial distribution. **(b)** Distribution of binding affinity measurements (*K*_D_) for all positive binders. **(c)** Identified hits for the challenging target TNF*α*. **(d)** Predicted structures for selected binder designs and corresponding *K*_D_ values for Chai-2 designed minibinders. Asterisk (^*^) indicates binding affinity measurements affected by avidity.

### 3.2 Double-digit success rates in *de novo* antibody design

Building on our early success with miniprotein design, we tackled the more challenging problem of antibody design. Computational antibody design has historically been more difficult than miniprotein design, likely due to the conformational flexibility of CDRs, the need to co-optimize two chains (heavy and light), and the mediation of binding by loops and beta-sheets, rather than the alpha helices common in previous miniprotein designs. As before, we prompt with a defined epitope on the target without any prior antibody structure or docking information. We also supply Chai-2 with a choice of antibody framework sequences among the five most common therapeutic VHH scaffolds and, separately, the five most common therapeutic VH-VL frameworks (extendable to additional scaffolds as needed). From these frameworks, Chai-2 designs all CDR residues (including their lengths) while preserving the chosen scaffold sequence. We apply this approach to the aforementioned broad, unbiased panel of 52 novel targets, spanning diverse protein families and subcellular localizations (Figure 3a), and test the designs experimentally in the wet-lab.

**Figure 3.**
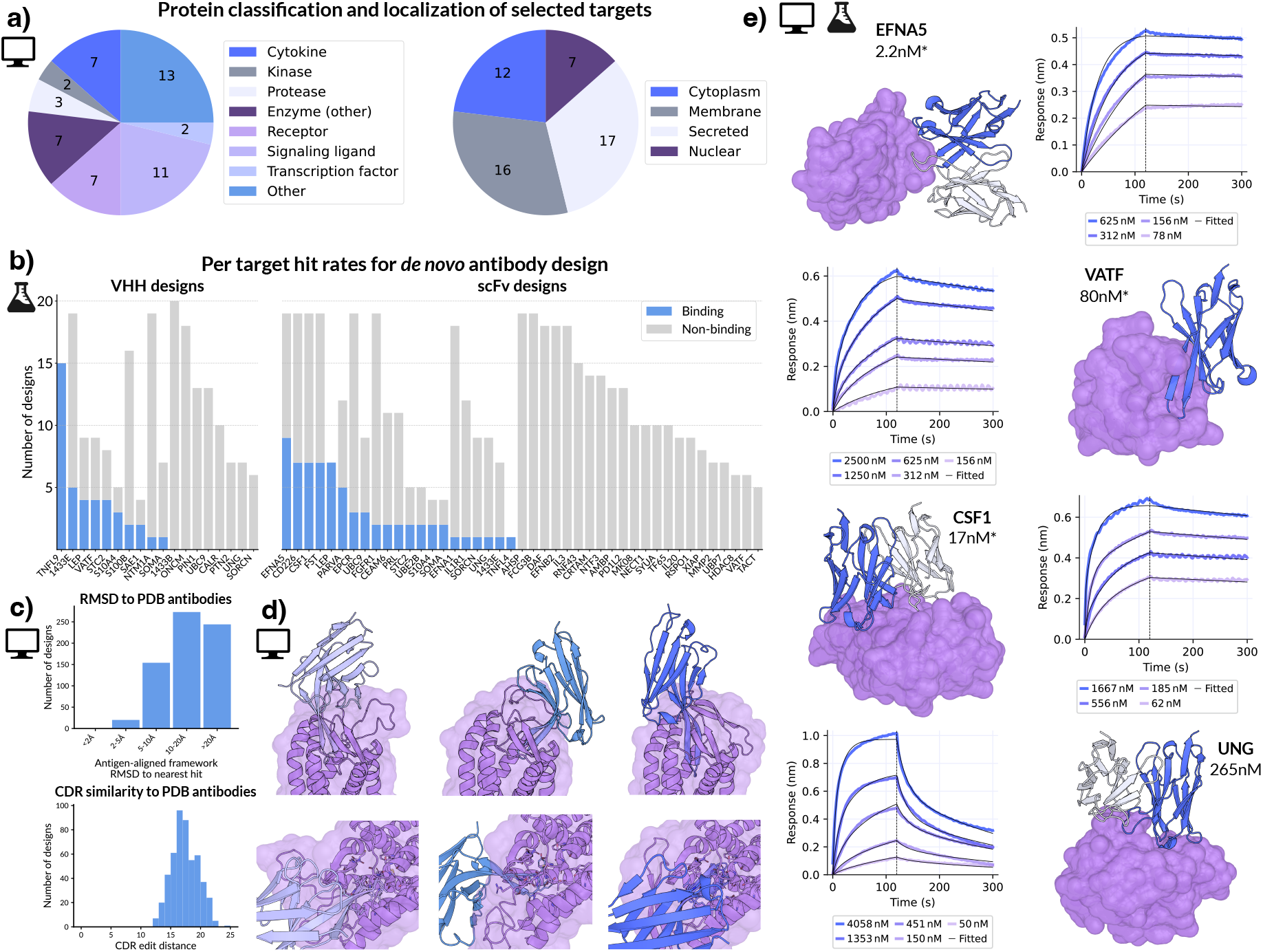
Antibody design targets and results. **(a)** Design targets annotated with protein classification (left) and subcellular localization (right). **(b)** Success rates for each target (x-axis), shown as the number of experimentally binding (purple) and non-binding (gray) *de novo* designs. **(c)** Structural (top) and sequence (bottom) novelty of experimentally tested designs with respect to known antibodies in the PDB. **(d)** Chai-2 generates diverse designs; top row shows three designed binders to one target, bottom row shows each binding interface in detail. **(e)** Examples of successful binder designs with BLI curves illustrating binding affinity. *K*_D_values marked with ^*^ are from avidity assay.

For each target and antibody format we select up to 20 designs for experimental validation (Figure 3b). Across the 52 total targets for which we have completed assays for scFv or VHH designs, we observe at least one binder for 26 of the targets, indicating that Chai-2 can create *de novo* antibodies for half of the proteins evaluated (Figure 3b). We observe an average hit rate of 15.5% across all designed antibodies (20.0% with VHH, 13.7% with scFv, Figure 1b and c) – an improvement of two orders of magnitude over prior state of the art [22]. We observe similar pass rates on a “hard” subset of these targets stringently filtered to remove even remote similarity (Figure S3), suggesting that these success rates reflect general design capabilities.

Although this evaluation explicitly focuses on targets with low similarity to chains with known antibodies, we nonetheless confirm that Chai-2 is not merely memorizing and retrieving the best training example during design. To do this, we used permissive sequence and structure searches to retrieve potentially similar antibody-antigen structures from the SAbDab [31] antibody database for each design, and calculated antigen-aligned framework RMSD (see Supplemental Information for details). The vast majority of our designs are at least 10Å RMSD from the most similar known antibody structure, and no designs are within 2Å of any existing antibody structure (Figure 3C, top). We also evaluate sequence similarity by computing the minimum CDR edit distance of each design to any antibody in SAbDab. All designs have a CDR edit distance *>* 10 from the closest example (Figure 3C, bottom). Together, these results indicate that the binders we generate are novel both by docking pose and by CDR sequences. We evaluate diversity within generated binders by clustering them by antigen-aligned antibody RMSD (Table S2). The majority of targets with at least two *de novo* binders contain multiple structural clusters, suggesting that Chai-2 explores multiple conformational states when designing successful binders. This range of structural diversity is qualitatively illustrated in Figure 3D.

In addition to exhibiting novelty, diversity, and high hit rates, our designed binders also exhibit high affinity. Figure 3e shows four examples of binders and their corresponding BLI curves. We assayed several of our binders against alternative targets to which they were **not designed** to bind to confirm that the high binding affinity is specific and not a consequence of indiscriminate binding (Table S3). We also experimentally screen for polyreactivity (Figure S4) and computationally screen our designs for developability and immunogenicity (Figure S5). These metrics are comparable to baselines, including existing therapeutic monoclonal antibodies, suggesting that beyond high hit rates, Chai-2 designs antibodies with many desirable properties amenable for further lead optimization and development.

### 3.3 Directly encoding design criteria in generation

Recognizing that therapeutic delivery requires more than just binder identification, we designed our pipeline for flexible prompting to address a range of design tasks. Below, we illustrate three key capabilities using targets that are not selected based on a training set holdout. We highlight flexible selection of antibody formats, precise epitope targeting, and direct design of antibodies with controllable cross-reactivity profiles.

To showcase format and epitope flexibility, we design antibodies to CCL2, a target involved in cardiovacular disease [32] and oncology [33, 34, 35]. From the same target protein, we generate VHH and scFv antibodies for two distinct epitopes. For each format–epitope pair, we experimentally characterize 20 designs. VHH designs yield 4 hits (20% hit rate) and scFv designs produce 5 hits (25% hit rate) (Figure 4a). While additional experiments are needed to confirm that the experimental binding sites match our *in silico* specifications, our results indicate that Chai-2 can successfully design binders in a variety of formats and follow fine-grained input prompts.

**Figure 4.**
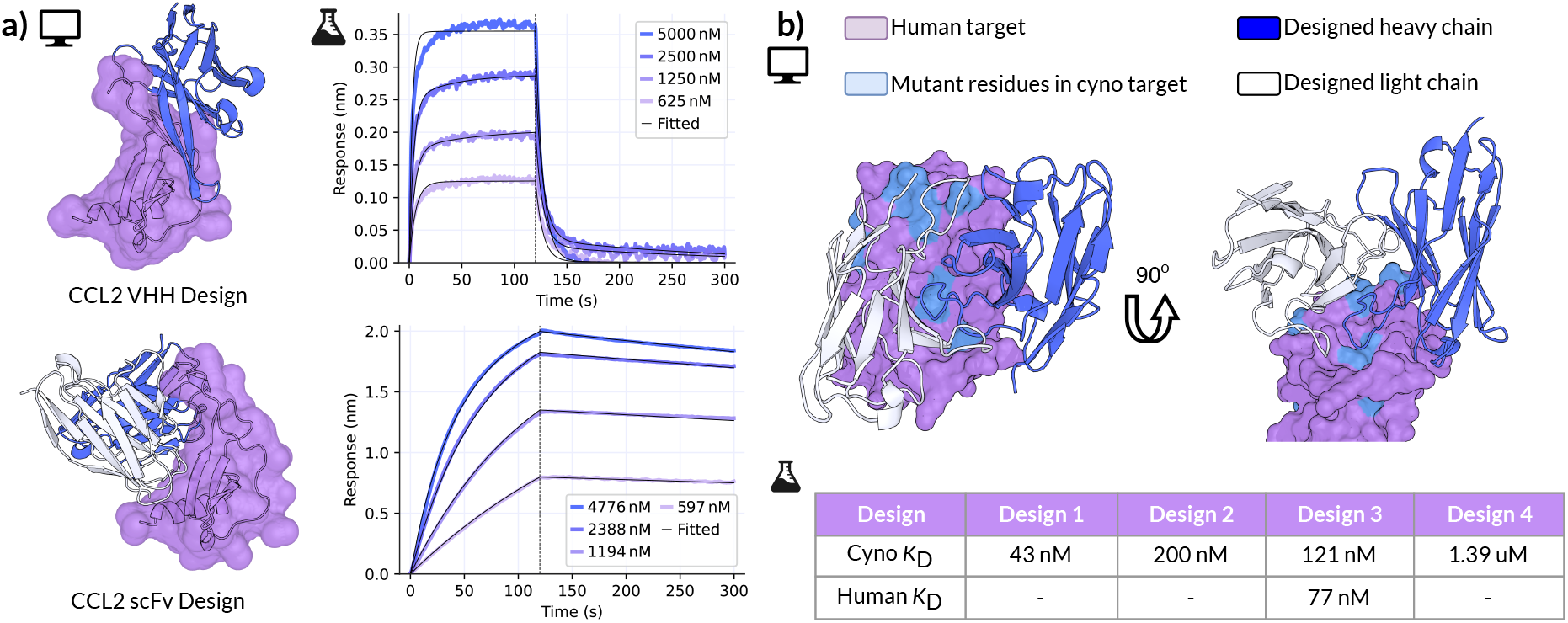
Chai-2 can be steered towards key antibody engineering tasks. **(a)** Our design platform can use different antibody formats (designed variable heavy in blue, designed variable light in white) when creating antibodies targeting different epitopes within the same target protein (purple). These two formats (VHH and scFv) both achieve double digit hit rates, and have best *K*_D_ values of 82 nM and 3.3 *μ*M, respectively (BLI curves, right). **(b)** Our design pipeline can also directly generate cross-reactive antibodies, shown here jointly optimizing human and cyno binding. We design heavy and light chains (blue and white, respectively) against the target (purple, cyno mutations in light blue). Both structures show the same design in different views. Of 14 designs selected for experimental profiling, we identify one hit with nanomolar affinities for both variants of the target (bottom table). Dashes indicate no detected binding.

Chai-2 can also design antibodies targeting multiple proteins in a single design run, enabling cross-reactive antibody engineering. This capability is particularly valuable for the development of single therapeutics that neutralize multiple variants of disease [36], or to streamline safety evaluation in preclinical studies using animal models [37]. As a case study, we prompted Chai-2 with homologous human and cyno sequences for a selected target, and advanced 14 candidates for experimental validation. We identified a lead antibody with dissociation constants of 77nM and 121nM against the human and cyno targets, respectively. Notably, not all binders had measurable binding to both homologs, suggesting that cross reactivity, in this case, is not trivially accomplished by targeting this epitope.

## 4 Future Work and Limitations

We have shown extensive lab validation for binding and specificity, and are currently investing in further characterization of our designs. To confirm our binders target the intended epitope, further competitive binding assays and laboratory 3D structure determination should be performed.

We note that all binding data collected are for scFvs and VHHs (see Section S5). The biophysical characteristics, such as affinities, of scFvs could differ from the same variable heavy and light chains reformatted as Fabs or full-length mAbs. Although scFv binding does not always translate to Fab binding, the formats can frequently be interconverted, and often times the steric constraints of Fabs even increase their affinities compared to scFvs of the same VH-VL sequences[38].

In this report, we primarily focus on binding characterization. We are actively working to characterize and improve the therapeutic properties of Chai-2 designs. For example, picomolar binders are frequently desirable for therapeutic applications [39, 40]. Furthermore, while we approximate humanness using *in silico* scores, further testing is required to fully understand potential immunogenicity liabilities. In a similar vein, biophysical assays measuring attributes such as thermal stability, aggregation propensity, and viscosity are required to validate that binders possess favorable profiles for downstream development. Future work will investigate more complex formats, such as bispecifics and antibody-drug conjugates, which make up a large component of biologics in development today.

The strong performance of Chai-2 in structure prediction – predicting 34% of antibody–antigen complexes with DockQ > 0.8 (compared to 17% for its predecessor, Chai-1) – highlights the power of integrating high-fidelity structure prediction with generative design. As structural accuracy continues to improve, we expect the fraction of targets tractable for computational binder design may grow proportionally. However, antibody CDR loops still present a notable challenge due to their intrinsic conformational flexibility, whereas the relative simplicity of modeling the *α*/*β* scaffolds of miniproteins likely underlie their comparatively higher hit rates and affinities. We suspect that design performance strongly depends on the underlying accuracy of structure predictions, as an incorrect atomic understanding of the problem can propagate into suboptimal choices in design.

## 5 Discussion

Chai-2 demonstrates state-of-the-art experimental success rates across diverse and challenging protein design tasks. Notably, the model achieves double-digit hit rates in wholly *de novo* antibody discovery across more than 50 targets and successfully generates high-affinity miniprotein binders against previously intractable targets such as TNF*α*. Remarkably, these outcomes are achieved while requiring orders-of-magnitude fewer physical measurements compared to existing approaches. Unlike traditional biologics discovery, which relies on extensive and often indiscriminate experimental screening, Chai-2 leverages a controllable, model-driven framework. By evaluating only 20 antibody or nanobody designs per target, we frequently identify strong binders within a two-week experimental cycle. Although higher affinities could likely be obtained by screening additional designs, our initial results indicate the potential to significantly compress discovery timelines—from months or years down to weeks.

The platform’s controllable and programmable design enables targeted exploration of molecular space, potentially unlocking modalities that elude existing techniques. In this work, we generate fully *de novo* designs in a single round while simultaneously optimizing epitope, scaffold, and specificity constraints. This shift from stochastic screening to intentional, programmable discovery suggests that antigens once deemed undruggable due to experimental challenges can potentially be addressed by *in silico* design. On-demand generation of epitope-specific binders could also streamline the development of advanced therapeutic formats such as antibody–drug conjugates, biparatopic constructs, and other multifunctional biologics. Furthermore, by reasoning at the atomic level—including ligands and post-translational modifications—Chai-2’s framework naturally extends beyond conventional biologics to macrocycles, peptides, enzymes, and small molecules. Collectively, we see our results as establishing computational-first design as an integral component of modern discovery platforms.

The hits designed by our model not only target epitopes without existing antibody designs, but also show high diversity in structure and sequence space. Looking ahead, we foresee a phased path to the generation of zero-shot drug candidates, where modeling of viscosity, pharmacokinetics, expression yield, and manufacturability could enable simultaneous optimization of multiple important attributes.

We believe that our results point toward a transition from empirical discovery to deterministic molecular engineering. By coupling atomic-resolution structure prediction with generative design, we short-circuit traditional discovery bottlenecks. Chai-2 marks a step toward the long-standing aspiration of rational drug design: computationally generating drug candidates that are ready for IND-enabling studies in a single shot, entirely on the computer.

## 6 Contributors

Jacques Boitreaud, Jack Dent, Danny Geisz, Matthew McPartlon, Joshua Meier, Zhuoran Qiao, Alex Rogozhnikov, Nathan Rollins, Paul Wollenhaupt, Kevin Wu.

## Supplemental Tables and Figures

**Figure S1.**
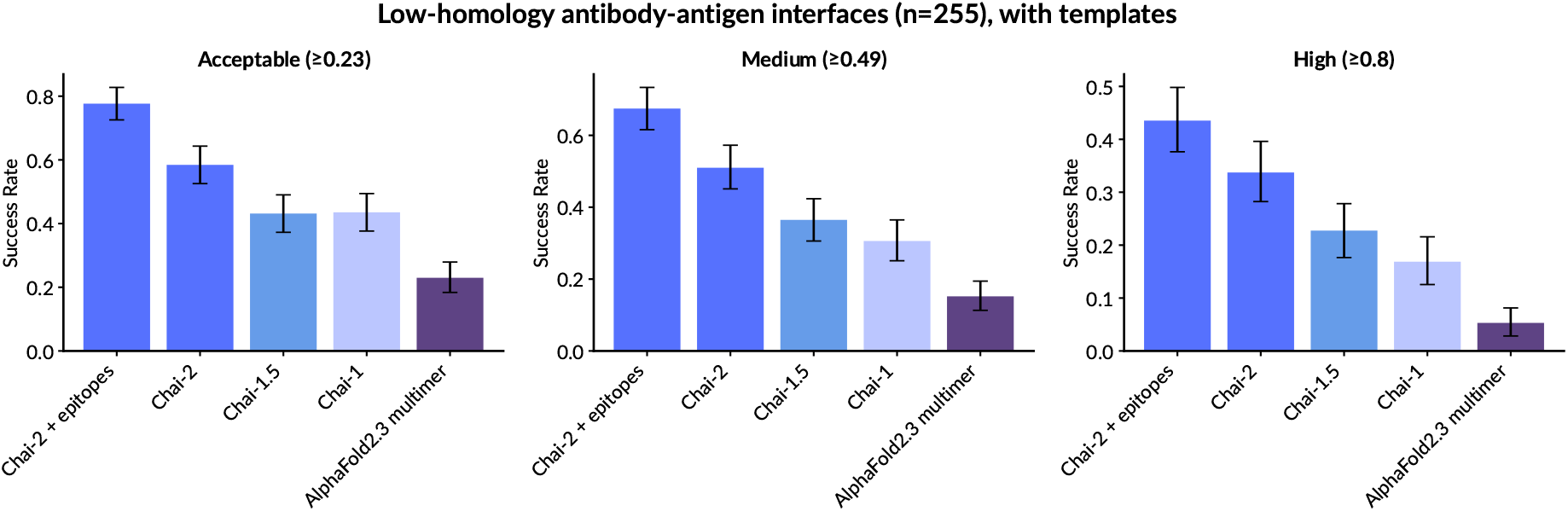
DockQ pass rates for low-homology antibody-antigen interactions, with templates. DockQ success rates are averaged across all interfaces. Error bars represent 95% confidence intervals from 10,000 bootstrap samples. Note that AlphaFold2.3 multimer sometimes uses templates that exactly match its input sequence thereby “leaking” the ground truth structure into model inputs, whereas all Chai models in this evaluation explicitly remove such templates to prevent data leakage. We consider high DockQ accuracy (*>* 0.8) to indicate predictions nearing experimental accuracy; see section S4. We cannot compare to AlphaFold3 due to its restrictions on commercial use. Chai-1.5 refers to an earlier version of the folding model, which was in development between the releases of Chai-1 and Chai-2. Chai-2 + epitopes refers to the Chai-2 model performance when prompted with four randomly selected antigen epitope resiudues using a 10Å C*α*-C*α* distance cutoff.

**Figure S2.**
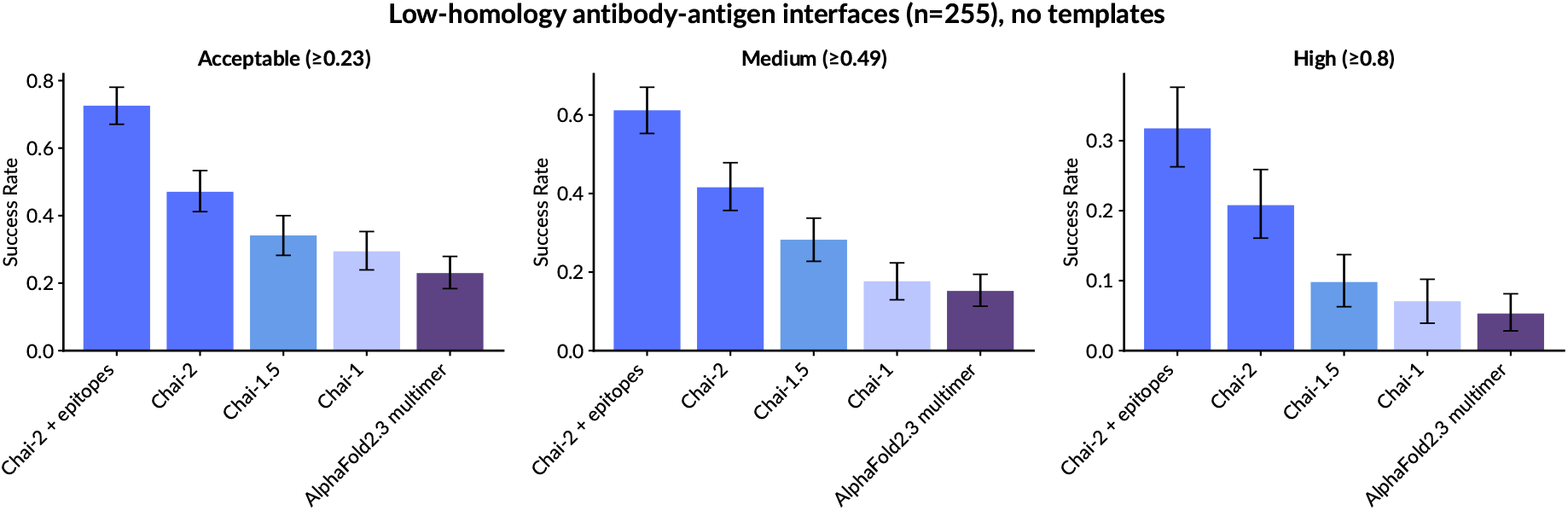
DockQ pass rates for low-homology antibody-antigen interfaces, without templates. DockQ success rates are averaged across all interfaces. Error bars represent 95% confidence intervals from 10,000 bootstrap samples. Templates provide solved structures of similar sequences; thus removing this information leads to a small but consistent performance dip compared to figure S1.

**Figure S3.**
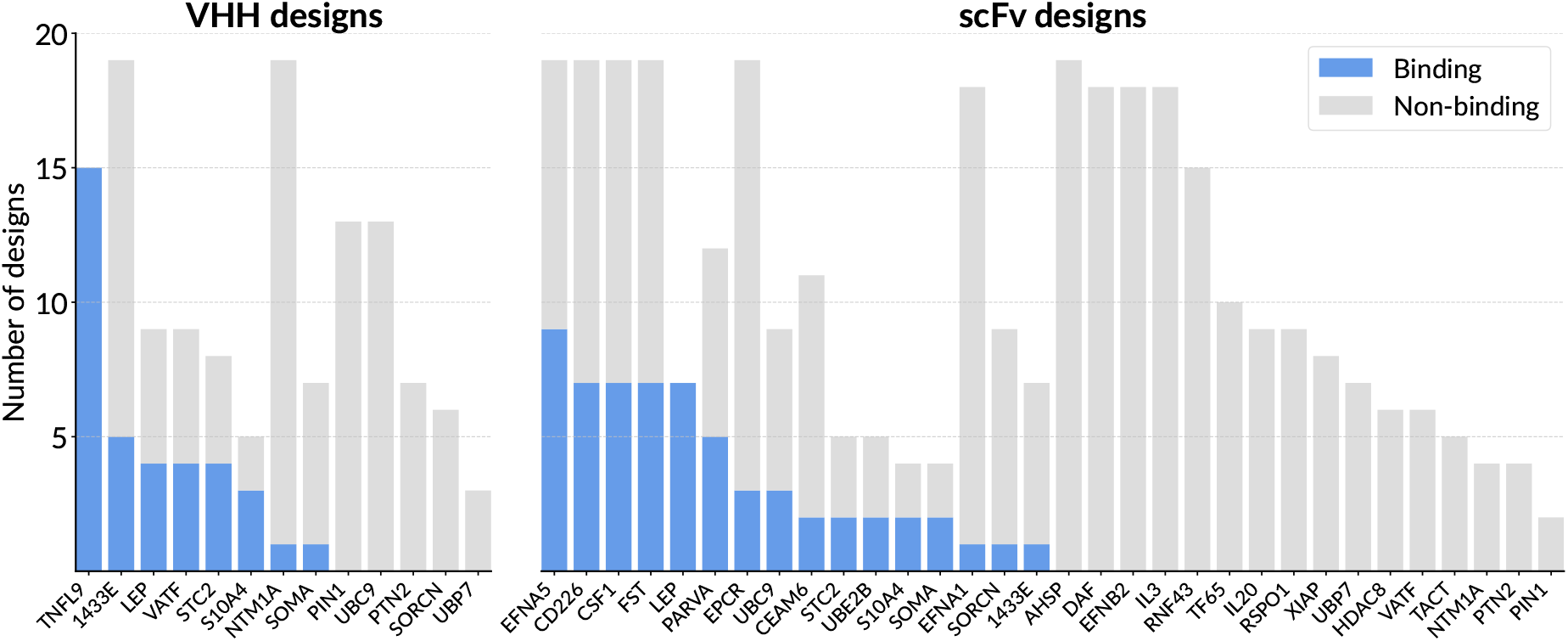
Per-target design pass rates for an ultra-low homology subset of targets. Compared to our main text, these targets are further filtered to remove targets with *>* 25% similarity and *>* 25% coverage to any SAbDab antigen prior to our training date cutoff. Of the 43 remaining targets, 24 (56%) have at least one binder (either for an scFv or VHH design); VHH designs have a binder success rate of 20% and scFv designs have a success rate of 17%. Despite the more challenging task, the success rates of Chai-2 in this setting closely match that of our full target panel.

**Table S1.** Design targets, their PDB entries, residue ranges, hotspot residues, binder lengths ranges used in Chai-2 prompts, and catalog item used for experimental validation. All chain IDs correspond to PDB subchain label.

| Design target | PDB ID | Target chain & residues | Hotspot residues | Binder length | Catalog item |
| --- | --- | --- | --- | --- | --- |
| IL-7RA [20, 28, 18] | 3di3 | B17-209 | B58, B80, B139 | 40-120 | Acro IL7-H52H7 |
| PD-L1 [20, 18] | 5o45 | A17-132 | A56, A115, A123 | 40-120 | Acro PD1-H5229 |
| InsulinR [20, 28, 18] | 4zxb | E6-155 | E64, E88, E96 | 40-120 | Acro INR-H52Ha |
| TNFα [20] | 1tnf | A12-157, B12-157, C12-157 | A113, C73 | 40-120 | Acro TNA-H4211 |
| PDGFRβ [28] | 3mjg | C93-280 | C265, C264, C210 | 50-120 | Acro PDB-H5259 |

**Figure S4.**
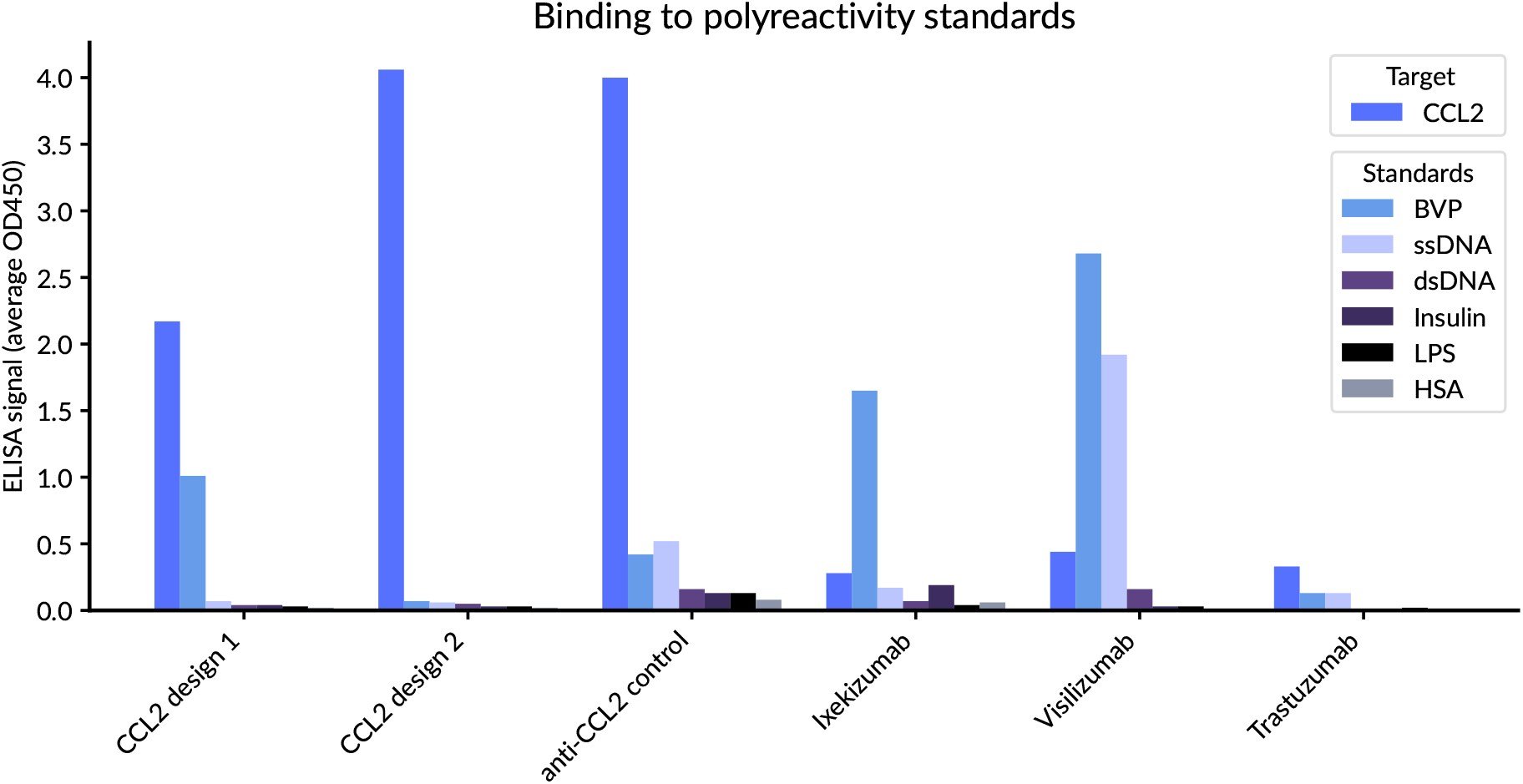
Experimental characterization of the polyreactivity of CCL2 designs to various standards. Ixekizumab and Visilizumab are examples of therapeutically approved, polyreactive mAbs, whereas Trastuzumab is an example of a “clean” therapeutic mAb. Our two *de novo* scFv designs against CCL2 exhibit comparable polyreactivity profiles to clinical therapeutics. The designs also show lower polyreactivity signal than a control anti-CCL2 antibody (11KP2 [41]) comparably reformatted as scFv. BVP: baculovirus particle; LPS: lipopolysaccharide; HSA: Human Serum Albumin. Further details can be found in Section S5.

**Figure S5.**
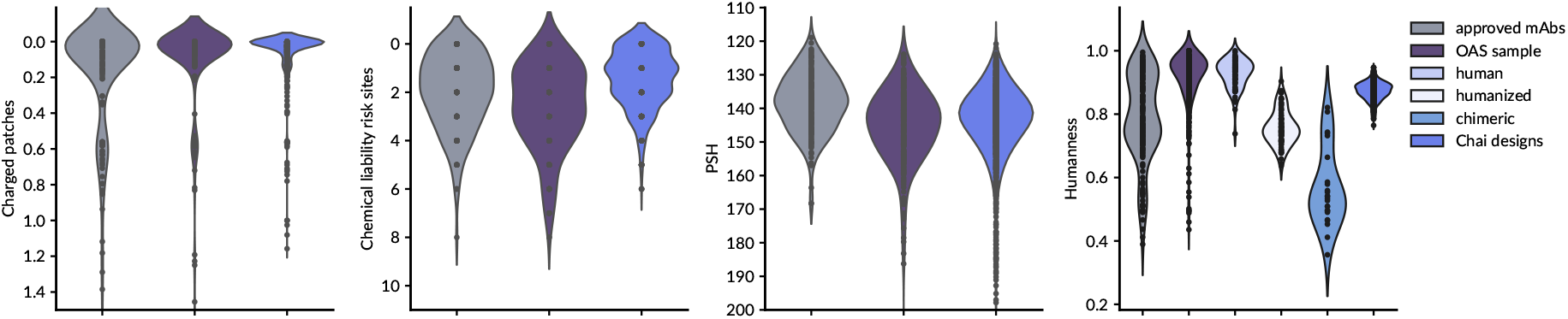
*In silico* immunogenicity and developability metrics. Compared to various baselines, Chai-2’s designs exhibit similar developability and immunogenicity profiles. We note that all designs are displayed in these plots, regardless of experimental binding determination. Exact computational protocols are defined in Section S3.3.

**Table S2.** Structural diversity of *de novo* antibody binders. A listing of all targets for which we have at least 2 experimentally confirmed antibody binders, as well as the number of structural clusters represented within those binders (binder-aligned antibody RMSD, clustered at 3Å). This clustering method primarily separates quaternary modes; there is additional diversity in precise CDR conformations that is not captured here. Of these 23 target-format combinations, 18 have two or more distinct structural clusters among their binding *de novo* antibodies.

| Format | Target name | # Structure clusters | # Hits | # Tested |
| --- | --- | --- | --- | --- |
| scFv | CD226 | 2 | 7 | 19 |
|  | CEAM6 | 1 | 2 | 11 |
|  | CSF1 | 2 | 7 | 19 |
|  | EFNA5 | 3 | 9 | 19 |
|  | EPCR | 3 | 3 | 19 |
|  | FGFR1 | 2 | 2 | 19 |
|  | FST | 3 | 7 | 19 |
|  | LEP | 2 | 7 | 7 |
|  | PARVA | 2 | 5 | 12 |
|  | PRL | 1 | 2 | 11 |
|  | S10A4 | 1 | 2 | 4 |
|  | SOMA | 1 | 2 | 4 |
|  | STC2 | 2 | 2 | 5 |
|  | UBC9 | 1 | 3 | 9 |
|  | UBE2B | 2 | 2 | 5 |
| VHH | 1433E | 3 | 5 | 19 |
|  | LEP | 2 | 4 | 9 |
|  | S100B | 2 | 2 | 16 |
|  | S10A4 | 2 | 3 | 5 |
|  | SAE1 | 2 | 2 | 4 |
|  | STC2 | 2 | 4 | 8 |
|  | TNFL9 | 4 | 15 | 15 |
|  | VATF | 2 | 4 | 9 |

**Table S3.**
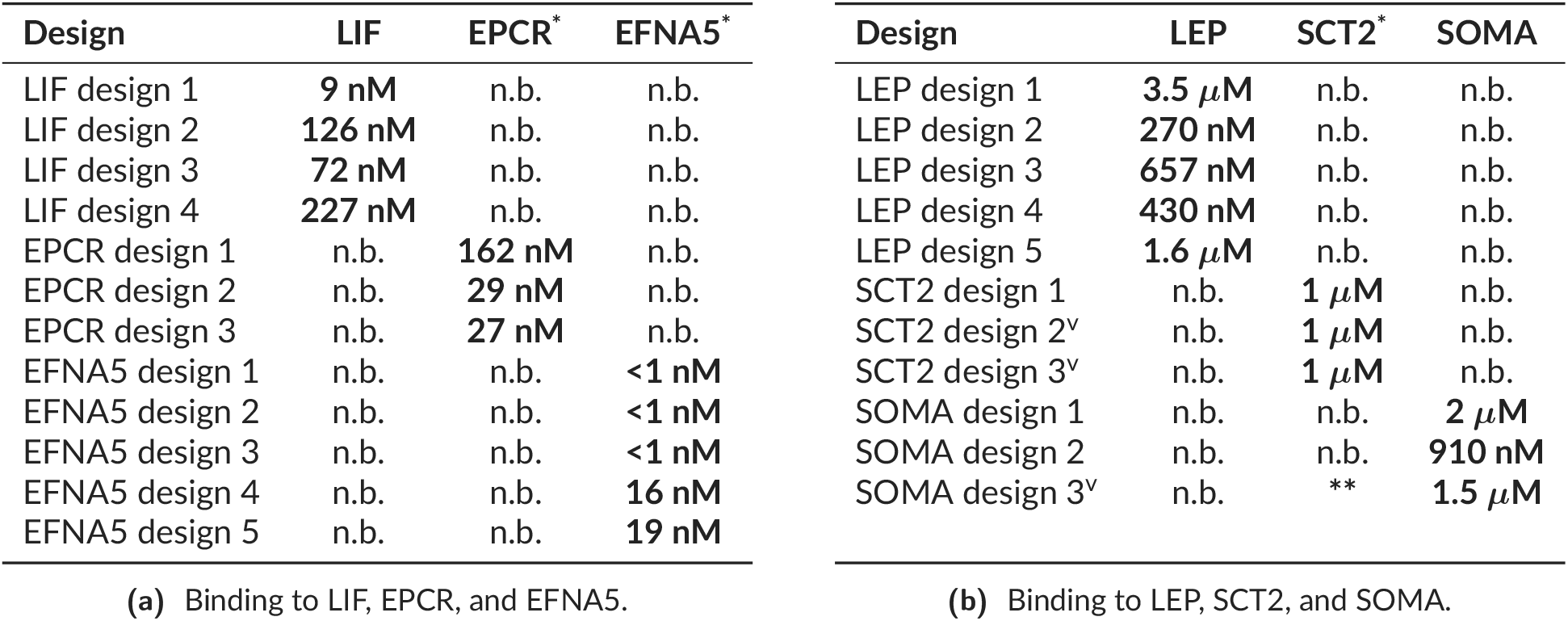
Most designs show no detectable binding to off-targets. We performed on-target, off-target BLI assays on two sets of our binding-positive *de novo* scFvs and VHHs: (a) designs targeting LIF, EPCR, EFNA5 and (b) designs targeting LEP, SCT2, and SOMA. In each experiment, all designs were measured for binding against all three antigens. Each on-target antigen reconfirmed positive binding signal and high affinity, whereas only 1 of the 23 designs (4%) showed binding above background to any off-target antigens. A designed VHH, SOMA design 3, showed binding above background to SCT2 but insufficient VHH material remained to determine the *K*_D_. Antigens were used as analyte at 5*μ*M concentration. The *de novo* scFv designs to LIF were excluded from the main benchmark as LIF has a known antibody in the PDB. (^*^) avidity measurement; (^**^) positive for binding, but no KD determined; (n.b.) no binding signal above background; (^v^) design is VHH-format, all other designs are scFv.

## Supplemental Information

### S1 Target selection

#### S1.1 Targets for miniprotein design

We use targets previously studied in Cao et al. [28], Watson et al. [18] and Zambaldi et al. [20], which focus on miniprotein design. We compare to experimental success rates reported in these papers. See Table S1 for exact design specifications used to produce results in Figure 2.

#### S1.2 Targets for *de novo* antibody design

We select a panel of protein targets for *de novo* antibody design from a catalog of proteins in stock in Contract Research Organization (CRO) catalogs, applying the following criteria:

- The protein should not have significant homology to any antigen sequence present in SAbDab [31] released prior to our models’ training date cutoff^2^, defined here as having at least 70% sequence identity covering at least 80% of the query (i.e., candidate target chain(s) for *de novo* design) as identified by `mmseqs`. By avoiding targets with high identity to solved antibody complexes, we reduce the confounding possibility that designs copy or adapt the structure or sequence of a known binder that might be cross-reactive. We also report results for a subset of these targets with no detectable similarity even at 25% identity with 25% coverage to ensure that our results are robust to targets very dissimilar to those seen during training.
- The antigen’s amino acid sequence should match exactly to some protein chain present in the PDB, where the chain is involved in a heteromeric interaction with another non-antibody protein chain.

For each selected target, we use an aforementioned heteromeric protein-protein binding interface to randomly sample epitope residues using a 10Å C*α*-C*α* distance as a cutoff. We provide a random subset of one to four of these residues to our model to guide sampling towards a specific interaction site. Notably, this selection procedure does not apply any heuristics that bias towards targets that may be more suitable for antibody binding, thus yielding a set of targets that should closely represent Chai-2’s capacity for generating binders to a broad, unbiased set of protein targets. We select 52 targets for this benchmark.

#### S1.3 Antigens with known antibodies

In this manuscript, we present results for antibody designs for two antigens with known antibodies in SAbdAb: CCL2 and LIF. These targets are used for profiling Chai-2’s ability to design antibodies with controllable format and epitope, for measuring cross-reactivity, and for assaying polyreactivity. We note that these targets are excluded from our calculation of *de novo* success rates.

### S2 Models and training

Chai-2 is a singular, end-to-end model for binder design. However, it incorporates several submodules that bear functional similarities to models used in the broader field of protein design. We describe two of these submodules: the design submodule Chai-2d, which is primarily responsible for proposing novel binders, and the folding submodule Chai-2f which is responsible for assessing the proposed designs from Chai-2d.

#### S2.1 Design submodule

The design submodule, Chai-2d, is trained to generate one or more protein chains that bind to arbitrary, pre-specified protein targets, and can be flexibly prompted to generate different “types” of binders. Chai-2d uses an all-atom generative protein model framework [21, 42] that simultaneously designs amino acid backbone and side chain atomic structure, extending these principles to binder design. Like the Chai-2f folding model, Chai-2d is not trained on structural data released after a temporal cutoff of 2021-9-30. Furthermore, Chai-2d is not trained on any structure having one or more protein chains with greater than 70% identity (and 80% coverage) to the targets reserved for design experiments. We identified such chains using `mmseqs easy-search` [43].

#### S2.2 Folding submodule architecture and training

Chai-2f follows a similar architecture as Chai-1 [12], similarly leveraging protein language model embeddings [44, 45, 46], multiple sequence alignments, and template information to predict structures. Like Chai-1, Chai-2f supports a variety of auxiliary restraint inputs that serve as additional conditioning information during the folding process.

Chai-2f is trained on synthetically predicted structures and the PDB [27]. PDB structures are subject to a date cutoff of 2021-9-30 (exclusive). We retain the data clustering and filtering strategies used to train Chai-1.

### S3 Metrics for designed antibody binders

#### S3.1 Sequence and structure novelty with respect to known antibodies

We evaluate the novelty of our designs by comparing to all antibody-antigen complexes in the PDB. For sequence novelty, we compute the minimum edit distance over concatenated CDRs between each design and all antibodies in the PDB. For structure novelty, we use a two step process where we first search for similar antigens in the PDB, then compare our designed structures to all retrieved antibody-antigen complexes. To maximize recall, we run antigen search using both structure and sequence based similarity. We use `foldseek easy-search` [47] with a TM score threshold of 0.5 and RMSD threshold of 5.0Å to find structurally similar matches, and `mmseqs easy-search` [43] with parameters `–min-seq-id 0.70 -c 0.25 –e-profile 1e-3 –cov-mode 1` to find similar matches in sequence space^3^. For each design, we compute the antigen-aligned alpha carbon (C*α*) RMSD of the heavy chain framework with respect to all retrieved structures, and report the RMSD of the closest hit. Alignment and RMSD values are computed with CEAlign [48] to handle variable length chains.

#### S3.2 Structure diversity within generated designs

Structural diversity of generated designs is computed within experimentally verified binders for a given target and of a given format. All such designs are aligned on the predicted folded structure of target chains, and we compute C*α* RMSD for all designed (antibody) chains. Note that unlike structural novelty, this computes RMSD across all residues and not just framework residues, such that the resulting distance metric includes conformational differences in CDR loop regions. This is done between each pair of designs, yielding a pairwise distance matrix that is then clustered with agglomerative clustering with mean linkage. Note that despite including CDRs in the calculation, the dominant contributor to target-aligned antibody RMSD is the ligand’s high-level placement and orientation relative to the target. Therefore this procedure primarily separates designs based on quaternary structure; there is additional structural diversity in fine-grained CDR loop conformations that is not captured by this calculation.

#### S3.3 Immunogenicity and Developability

Humanness of designs was evaluated using the `promb` package (https://github.com/MSDLLCpapers/promb) following methodology proposed in BioPhi [49]. Chemical liability risk sites were identified following the approach of Satława et al. [50], with two modifications: only high-severity liabilities were included, and regions containing three consecutive hydrophobic residues were also classified as liabilities. Hydrophobic patches (PSH) and charged patches score (sum of patches of positive/negative charge) were implemented according to Raybould et al. [51].

To establish a baseline for which we can compare our designs, we consider the following controls: approved mAbs from Thera-SAbDab [52], randomly sampled antibodies from Paired OAS [53], and different groups from the “217 Immunogenicity” dataset from Prihoda et al. [49] (originally curated in Marks et al. [54]).

**Table S4.** Folding model inference settings. All other settings are kept at default values. We run multiple AlphaFold 2.3 models, each with different seeds to mimic the trunk and diffusion sampling strategy more common in recent folding models. We use 3 recycles according to the ColabFold implementation, and do not expect a significant change if more are used.

| Model | Trunk samples | Diffusion samples | Models | Seeds | Recycles |
| --- | --- | --- | --- | --- | --- |
| Chai-1 | 5 | 5 | - | - | 3 |
| Chai-2f / Chai-1.5f | 5 | 5 | - | - | 10 |
| AlphaFold 2.3 multimer | - | - | 5 | 5 | 3 |

### S4 Folding model evaluation on antibody-antigen complexes

We benchmark the following folding models, all of which were the latest versions available as of June 17, 2025.

- **Chai-1** [12] https://github.com/chaidiscovery/chai-lab, commit SHA `103dc24`
- **AlphaFold 2.3 multimer via ColabFold** [55, 56] https://github.com/sokrypton/ColabFold, using the authors’ pre-built docker image at ghcr.io/sokrypton/colabfold:1.5.5-cuda12.2.2

All Chai models were provided with the same MSAs generated by `mmseqs easy-search` run against `uniref2302` and `colabfold_envdb_202108` databases as provided by the ColabFold project https://colabfold.mmseqs.com [56]. When provided, templates are obtained running `mmseqs easy-search` against the `pdb100_230517` database, and are subject to a 95% sequence identity filter to exclude exact hits. AlphaFold2.3 was run via Colabfold [56] with server-provided MSAs and templates (when enabled). During this evaluation, we found that Colabfold’s server templates can sometimes contain an exact match for the protein chain being folded and so AF2.3 can have access to additional information that Chai models do not have. We are unable to benchmark AlphaFold3 due to licensing restrictions. All models were run with settings listed in table S4.

We adopt DockQ [57, 58] as our primary evaluation metric, using the official implementation (https://github.com/bjornwallner/DockQ, version 2.1.1). We fold each complex with all chains present, then calculate DockQ specifically between the low homology interfaces in our evaluation set (see below). We follow standard values for thresholding DockQ values into successful (*≥*0.23), medium (*≥* 0.49), and high (*≥* 0.80) quality predictions. We consider performance in the DockQ high range to indicate predictions near experimental accuracy, these generally require predictions to have sub-angstrom RMSD accuracy at the binding interface.

We conduct an antibody-focused evaluation of folding models by using the low-homology evaluation set previously presented in Chai-1 [12], but keeping only examples that overlap SAbDab [31], contain an antibody-antigen interface as annotated by ANARCI [59], do not contain DNA or RNA chains (for comparability to AlphaFold 2.3 multimer), and do not exceed 1024 tokens in length. This yields 152 complexes with 255 low homology interfaces, with release dates spanning 2022-05-11 to 2023-01-11. On these low homology antibody antigen interfaces, we find that our folding models have consistently and meaningfully improved on this difficult task. Compared to Chai-1, which was released less than a year ago, Chai-2 doubles the rate at which interfaces are predicted at experimental accuracy (i.e., DockQ exceeding 0.8, Figure S1). The performance delta increases to three-fold when templates are not provided to either Chai or AlphaFold models (Figure S2). We further note that when epitope restraints are provided – as often is the case for design, as we know the epitope *a priori* – there is a dramatic increase in DockQ success rates, with the rate of structures predicted with experimental accuracy increasing to above 40% with templates, or 32% without.

### S5 Experimental Methods

#### S5.1 Protein production

Each design sequence has a C-terminal purification tag added and was reverse-translated to DNA sequence by a codon optimization algorithm for high production in *E. coli*. The sequence was synthesized as gene fragments, cloned into expression vector, and expressed in an E. coli-based *in vitro* transcription–translation system.

##### Minibinders

Miniproteins used a Twin-Strep tag and were expressed in 8*μ*L volume at 37°C for 12 hours. Miniprotein concentration was quantified using a tag-specific fluorescence assay, each concentration was normalized, and the protein was purified by direct capture and washing on the streptavidin-coated BLI probe.

##### Antibodies and nanobodies

scFvs and VHHs used 10xHis tag and were expressed in 1mL volume at 30°C for 3 hours. The reactions were centrifuged at 18,000 x g and 4 °C for 10min. Supernatent was collected and purified by Ni column. Concentrations were determined by OD280 measurement and purity was confirmed by Coomassie blue-stained SDS-PAGE gel under reducing conditions.

#### S5.2 Binding by Bio-Layer Interferometry (BLI)

##### Minibinders

Multi-cycle kinetic assays were conducted by BLI (Gator Bio). The designs were immobilized on streptavidin-coated probes via the Twin-Strep tag. Target protein solutions at 1000, 316.2, 100, 32.2, and 0 nM were flowed over the probes, with signal acquisition at a sampling rate of 5 Hz. The BLI assay sequence was as follows: 120s baseline, 120s ligand loading, 200s post-loading baseline, 220s association phase, 240s dissociation phase, and 75s regeneration. Measurements were made at 25°C with 50 mM HEPES, 100 mM NaCl, and 0.5% Triton X-100 as kinetic buffer. Negative controls, including buffer-only samples, were included for baseline subtraction. Data was fit globally to a 1:1 binding model across all concentrations to obtain affinity constants (*K*_D_).

##### Antibodies and nanobodies

Multi-cycle kinetic assays were conducted by BLI (Octet RED384). Generally, the designed scFvs / VHHs were immobilized on HIS1K (anti-penta-HIS) probes via the His tag, and target was applied as analyte. A subset of targets (1433B, CALR, CEACAM-6, CRTAM, FCGR3B, NTF3, ONCM, PRL, S100B, and UNG) were instead immobilized on SA sensor and designs were analyte. For the binding hit screen, 1-point BLI was performed with either 5*μ*M or 10*μ*M analyte. For *K*_D_determination 4- to 7-point series of 2- or 3-fold dilutions were performed. Signal acquisition used a sampling rate of 5 Hz. The BLI assay sequence was as follows: 60s baseline, 300s ligand loading, 100s post-loading baseline, 120s association phase, 180s dissociation phase, and 30s regeneration. As necessary the association and dissociation phases were extended up to a maximum of 300s each to allow for adequate saturation or dissociation. Measurements were made at 25°C, with 10 mM Na2HPO4 12H2O, 2 mM KH2PO4, 137 mM NaCl, 2.7 mM KCl, 0.05% Tween-20, 0.1%BSA pH7.4 (PBST+0.1%BSA) as kinetic buffer. Negative controls, including buffer-only samples, were included for baseline subtraction. Data was fit globally to either a 1:1 or 1:2 binding model across all concentrations to obtain *K*_D_.

Across all formats, designs with binding-positive curve signature, signal greater than 300% of the negative background signal, and signal greater than 0.1nm above background were classified as positive hits. *K*_D_values in this report were all obtained from fits with R2 > 0.98. Positive controls with known affinity were included for all targets for assay validation. A list of all BLI target materials can be found in Table S5.

#### S5.3 Polyreactivity assays

ELISA for polyreactivity to BVP was performed as described in Jain et al. [60]. Sample concentration was 3.5*μ*M for scFvs and 1*μ*M for Ixekizumab (MedChem #HY-P9924), Visilizumab (MedChem #HY-P99332), and Trastuzumab (MedChem #HY-P9907). ELISA for polyreactivity to ssDNA, dsDNA, insulin, LPS and HSA was performed as described in Mouquet et al. [61]. ssDNA (Sigma #D8899), dsDNA (Sigma #D4522), insulin (Yeasen #40112ES80), LPS (Yeasen #60747ES08) and HSA (Sino #10968-HNAY) were coated at 1*μ*g/mL. Sample concentration was 10*μ*g/mL for all test articles. In both assays, 0.15 *μ*g/mL anti-His/HRP was used to detect His-tagged scFv and 0.08 *μ*g/mL anti-huIgG/HRP was used to detect control antibodies. Plotted OD450 values are the average of two replicates with blank subtracted.

**Table S5.** Target material for each antigen used in wet-lab experiments.

| Target | Catalog item |
| --- | --- |
| 1433B | Sino 10843-H09E-B |
| 1433E | Sino 50691-M09E |
| AHSP | Sino 14391-HNAE |
| AMBP | Sino 13141-H05H1 |
| CALR | Sino 13539-H02H |
| CD226 | Sino 10565-H02H |
| CEAM6 | Sino 10823-H08H-B |
| CRTAM | Sino 11975-H08H-B |
| CSF1 | Sino 11792-H02H |
| DAF | Sino 10101-H02H |
| EFNA1 | Sino 10882-H02H |
| EFNA5 | Sino 10192-H02H |
| EFNB2 | Sino 10881-HCCH |
| EPCR | Sino 13320-H02H |
| FCG3B | Sino 11046-H27H-B |
| FGFR1 | Sino 10616-H02H |
| FST | Sino 10685-H02H |
| HDAC8 | Sino 10864-H09B |
| IL1R1 | Sino 10126-H02H |
| IL20 | Sino 13060-HNAE |
| IL3 | Sino 11858-HNAE |
| LEP | Sino 10221-HNAE |
| MK08 | Sino 10795-H09B |
| MMP2 | Sino 10082-HNAH |
| NECT1 | Sino 11611-H02H |
| NTF3 | Sino 10286-HNAE-B |
| NTM1A | Sino 11222-HNCE |
| ONCM | Sino 50112-MNAE-B |
| PARVA | Sino 13919-H09E |
| PD1L2 | Sino 10292-H02H |
| PIN1 | Sino 10282-HNCE |
| PRL | Sino 10275-HNAE-B |
| PTN2 | Sino 10570-HNCB |
| RNF43 | Sino 16108-H02H |
| RSPO1 | Sino 11083-HNAS |
| S100B | Sino 10181-H01H-B |
| S10A4 | Sino 10185-H01H |
| SAE1 | Sino 13921-HNCB |
| SOMA | Sino 16122-HNCE |
| SORCN | Sino 14547-HNCB |
| STC2 | Sino 13653-H02H |
| SYUA | Sino 12093-HNAE |
| TACT | Sino 11202-H08H-B |
| TF65 | Sino 12054-H09E |
| TNFL4 | Sino 13127-H04H |
| TNFL9 | Sino 15693-H01H |
| UBC9 | Sino 13205-HNCE |
| UBE2B | Sino 51243-MNCE |
| UBP7 | Sino 11681-HNCB |
| UNG | Sino 12939-H09E-B |
| VATF | Sino 15605-H09E |
| XIAP | Sino 10606-H17E |

## Footnotes

1 For convenience, we test slightly different numbers of designs per miniprotein target.

2 SAbDab identifiers referencing obsolete PDB identifiers are assumed to be have been released prior to training date cutoff for simplicity and caution.

3 This uses a more permissive coverage cutoff of 25% with respect to the target compared to how we identified *de novo* targets.

